# Beyond power limits: the kinetic energy capacity of skeletal muscle

**DOI:** 10.1101/2024.03.02.583090

**Authors:** David Labonte, Natalie C Holt

## Abstract

Muscle is the universal agent of animal movement, and limits to muscle performance are therefore an integral aspect of animal behaviour, ecology, and evolution. A mechanical perspective on movement makes it amenable to analysis from first principles, and so brings the seeming certitude of simple physical laws to the challenging comparative study of complex biological systems. Early contributions on movement biomechanics considered muscle energy output to be limited by muscle work capacity, *W*_max_; triggered by seminal work in the late 1960s, it is now held broadly that a complete analysis of muscle energy output is to also consider muscle power capacity, for no unit of work can be delivered in arbitrarily brief time. Here, we adopt a critical stance towards this paradigmatic notion of a power-limit, and argue that the alternative constraint to muscle energy output is instead imposed by a characteristic kinetic energy capacity, *K*_max_, dictated by the maximum speed with which the actuating muscle can shorten. The two critical energies can now be directly compared, and define the physiological similarity index, Γ = *K*_max_*/W*_max_. It is the explanatory power of this comparison that lends weight to a shift in perspective from muscle power to kinetic energy capacity, as is argued through a series of brief illustrative examples. Γ emerges as an important dimensionless number in musculoskeletal dynamics, and sparks novel hypotheses on functional adaptations in musculoskeletal “design” that depart from the parsimonious evolutionary null hypothesis of geometric similarity.

---

Movement is essential for all animals, and muscle is what drives animal movement (Alexander, 2003; Biewener and Patek, 2018; Biewener, 2016; Daniel and Tu, 1999; Dickinson et al., 2000; Higham et al., 2016; McMahon, 1984; Mendoza et al., 2023). What muscle can and cannot do is thus a fundamental question in zoology. Two rudimentary mechanical properties are thought to characterise each unit of muscle mass as a motor: its maximum work density, and its maximum power density (Bennet-Clark, 1977; Borelli, 1680; Gabriel, 1984; Hill, 1950b). No muscle contraction can violate these limits, and, because both muscle power and work density appear to be remarkably conserved (they vary by at most one order of magnitude across animal size, ecological niche, and evolutionary history, see Askew and Marsh, 2002; Close, 1972; Marden and Allen, 2002; Medler, 2002; Rospars and Meyer-Vernet, 2016), they are thought to pose universal constraints on animal performance. A limiting work or power density have, in some form or another, been invoked to account for a remarkable diversity of non-trivial observations on animal locomotor performance, including the maximum running speed (e. g. Hill, 1950b; Irschick et al., 2003; Labonte et al., 2024; Meyer-Vernet and Rospars, 2015; Usherwood and Gladman, 2020), flight speed (Alexander, 2005; Pennycuick, 1968), swimming speed (e. g. Clemente and Federle, 2012; O’dor and Webber, 1991; Richards and Clemente, 2013; Richards and Sawicki, 2012; Wakeling and Johnston, 1998), and jump height (e. g. Bennet-Clark, 1977; Gabriel, 1984; Lutz and Rome, 1994; Marsh, 1994; Roberts et al., 2011; Sutton et al., 2016; Wilson and James, 2000); size-specific variations in animal posture (Usherwood, 2013); the prevalence of latched “power-amplifiers” in small animals (Alexander, 1988; Bennet-Clark, 1975; Gronenberg, 1996; Ilton et al., 2018; Longo et al., 2019; Patek, 2023); the transition from latch-mediated to direct muscle actuation in jumping animals (Sutton et al., 2019); the mechanical benefits of in-series elasticity in explosive muscle contractions (e. g. Aerts, 1997; Alexander, 1995; Galantis and Woledge, 2003; Hill, 1950a; James et al., 2007; Lichtwark and Wilson, 2005; Mendoza and Azizi, 2021; Peplowski and Marsh, 1997; Roberts, 2016; Roberts and Azizi, 2011; Roberts and Marsh, 2003; Rosario et al., 2016; Sawicki et al., 2015, reviewed recently in Holt and Mayfield (2023)); the outcome of predator-prey interactions (Wilson et al., 2018); feeding performance (Camp et al., 2015); or the limits to manoeuvrability (Williams et al., 2009; Wilson et al., 2013), to name but a few examples.

The notion of a work- and power-limit on muscle energy output now pervades the comparative biomechanics literature, but it originated in the analysis of animal jump performance (Bennet-Clark, 1977); the concept is thus perhaps best introduced through this historical lens.

## Two laws to bind jump height

Animal jump performance is thought to be bound by two “laws”: Borelli’s law, which encodes a work constraint to muscle energy output (Borelli, 1680); and what arguably should be called Bennet-Clark’s law, which prescribes a power constraint (Bennet-Clark, 1977; Bennet-Clark and Lucey, 1967).

The argument for the work constraint typically runs as follows (Alexander, 2003; Biewener and Patek, 2018; Hill, 1950b; McMahon, 1984; Schmidt-Nielsen, 1984): an animal that jumped to a height *h* has increased the gravitational potential energy of its centre-of-mass by an amount *E*_*pot*_ = *mgh*. This increase was paid for with mechanical work, done by muscle at the expense of chemical energy.

The maximum mechanical work muscle can do depends on its volume, *V*, and on the maximum displacement-averaged stress, 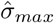, it exerts as it shortens by a maximum fraction of its length, *ε*_*max*_,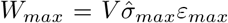. An estimate for the maximum jump height then follows via conservation of energy, *E*_*pot*_ = *W*_*max*_, where it is tacit that all external forces are much smaller than the muscle force (Scholz et al., 2006, see SI for a more detailed discussion of this point.). The physiological parameters 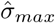, *ε*_*max*_, and the muscle volume per unit body mass, *V/m*, are typically considered size-invariant; animals small or large should consequently be able to deliver the same mass-specific energy with their muscle, and thus jump to the same height, *h*_*max*_ = *V σ*_max_*ε*_max_(*mg*)^−1^. It is this prediction that is often referred to as *Borelli’s law*.

The presentation of the power constraint usually remains on curiously more qualitative grounds. Bennet-Clark himself writes: “As the time available for acceleration is less in smaller animals, the energy store must be able to deliver the energy more rapidly” (Bennet-Clark, 1977). Schmidt-Nielsen explains (Schmidt-Nielsen, 1984): “The smaller the animal, the shorter its take-off distance. […] The time available for take-off is very short, and muscle just cannot contract that fast”. Alexander has it that “smaller jumping animals have smaller acceleration distances, and so have to extend their legs in even shorter times […]. But no known muscle can complete an isolated contraction in so short a time.” (Alexander, 2003). Biewener and Patek agree (Biewener and Patek, 2018): “Smaller animals have shorter limbs […]; therefore, the […] time available for acceleration during take-off is less”. These representative accounts of how power may constrain muscle energy output have two elements in common. First, they all highlight the importance of time. Time is absent in the framework of Borelli’s law, because it frames the problem solely in terms of muscle work. But no fixed amount of work can be done in an infinitely short amount of time, and time therefore ought to be considered explicitly when estimating bounds on muscle mechanical performance. Second, it is implied that the time available depends on animal size. Smaller animals are thought to have less time to do work, and muscle energy output, so the argument goes, is thus constrained by muscle power capacity in animals below some critical size, but by muscle work capacity in animals above it (e. g. Alexander, 2003; Bennet-Clark, 1977; Biewener and Patek, 2018; Gabriel, 1984; Ilton et al., 2018; Longo et al., 2019; Patek, 2023; Schmidt-Nielsen, 1984; Usherwood, 2013; Usherwood and Gladman, 2020). It is the omission of time that prompted Bennet-Clark and others to challenge Borelli’s law; and it is the suggested variation of available time with animal size that eventually developed into the now practically ubiquitous notion of a power-limit to the energy output of muscle—Bennet-Clark’s law.

This manuscript revisits this power-limit paradigm, and presents an alternative mechanical framework to account for size-specific variations in muscle energy output.

## Preliminaries, model formulation and assumptions

To formally assess the mechanical grounds on which the notion of a power-limit to muscle energy output rests, it is prudent to first lay out what is meant by it. It is obviously correct that an isolated muscle with mass *m*_*m*_ and power density *P*_*ρ*_ cannot provide more power than *P*_max_ = *P*_*ρ*_*m*_*m*_. But this truism merely provides the basis for a more complex and consequential causal inference widespread in the comparative biomechanics literature: the power-limit is regularly invoked as a constraint on the *energy* muscle can provide with each contraction; it is branded as the proximate cause of an ultimate limit distinct from Borelli’s law. This assertion has two corollaries: changes in muscle power must be necessary and sufficient to change muscle energy output, for otherwise it is unclear in what sense power can be said to be limiting; and the changes in power that lead to a variation in energy output must leave muscle work capacity unaffected, for otherwise there is no clear basis upon which Borelli’s and Bennet-Clark’s law may be argued to be different. These two demands define the burden of proof that rests with any claim of an energy limit distinct from Borelli’s law; with them at hand, an assessment strategy can be formulated.

To probe the validity of the first corollary, we will analyse how much energy a muscle with finite work and power capacity can deliver in a single contraction, using the mathematically simplest form of physical reasoning— dimensional analysis. The aim here, as much as for the rest of this study, is not to account for all complexities of real musculoskeletal systems, but to analyse parsimonious models that capture the essential physical features that endow a muscle with a maximum work and power capacity—the properties that underpin the claims under investigation. A suitable framework for such an analysis can be found in the recently developed theory of physiological similarity, which maps out the mechanical performance landscape for an idealised musculoskeletal system (Labonte, 2023; Labonte et al., 2024; Polet and Labonte, 2024). This system consists of a payload of mass *m*—for example the body mass of an organism, or the mass of a limb that is moved—connected to a muscle of volume *V*, which can exert a maximum stress *σ*_max_, shorten by no more than a maximum strain *ε*_max_, and no faster than with a maximum strain rate 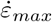. Throughout this work, it is assumed that the muscle is fully activated throughout the contraction, and that the payload mass is large compared to the muscle mass, *m >> m*_*m*_.

Without loss of generality, such a muscle has a maximum work capacity *W*_max_ ∝ *V σ*_max_*ε*_max_. The muscle’s power capacity, in turn, may be defined in one of two ways. Early work on power-limits in small animals usually refers to the *time-averaged power* —the ratio between the work done and the time it took to deliver it, 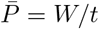; this focus on 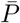 is often implicit, identifiable only from reference to the time available. More recent work also considers the instantaneous power, *P*_inst_, the product between the instantaneous force, *F*, and instantaneous velocity, *v, P*_inst_ = *W/dt* = *Fv*. The time-averaged power is easier to measure, and is arguably the more functionally relevant metric; but the limit on instantaneous muscle power is mechanically and physiologically important, and muscle cannot exceed either. Both powers are proportional to muscle volume, stress and strain rate, 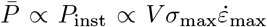, so that the distinction is irrelevant at this point, and no further restrictions need be imposed.

To analyse the physical limits to muscle mechanical performance with respect to the second corollary, we will then deploy the fundamental principle of the conservation of energy. This step, still conducted within the framework of the theory of physiological similarity, will force as much as permit us to take into account a key feature of real muscle that is irrelevant for the results of the initial dimensional analysis, and was thus ignored up to now: muscle stress typically varies inversely with muscle strain rate according to the well characterized force-velocity relationship (FVR; Hill, 1938; Piazzesi et al., 2007), 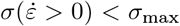 (for a model that also considers force-length properties, see Labonte, 2023). This co-variation of stress with strain rate stems directly from the molecular mechanisms that underpin muscle contraction (Hill, 1938; Piazzesi et al., 2007), and places further constraints on muscle energy output, for the muscle’s work capacity now declines with the rate at which it is delivered (Bobbert, 2013; Peplowski and Marsh, 1997; Roberts and Marsh, 2003). The task ahead is thus not only to formally identify all possible limits on muscle energy output, but also to evaluate explicitly how the variation of stress with strain rate influences how much energy muscle can inject in one single contraction.

Three important features that characterise real musculoskeletal systems are notably absent from the idealised model under investigation, and thus demand commentary. First, muscle input is usually proportional but not equal to system output: muscle input is transmitted through joints, resulting in musculoskeletal systems that are geared (Borelli, 1680; Osborn, 1900). Gearing leaves the system’s maximum work- and power-capacity unaffected, and is thus only discussed briefly; readers who seek a detailed treatment of the influence of gearing on muscle energy and power output may find it in Arnold et al. (2011); Labonte (2023); McHenry (2011, 2012); Polet and Labonte (2024). Second, muscle rarely attaches directly to skeletal segments, but instead connects onto them via elastic elements such as tendons, aponeuroses and apodemes (Hill, 1950a). The resulting in-series elasticity plays an important role in the power- and energy-output of skeletal muscle (Aerts, 1997; Alexander, 1995; Bennet-Clark, 1975; Galantis and Woledge, 2003; Hill, 1950a; Holt and Mayfield, 2023; James et al., 2007; Lichtwark and Wilson, 2005; Mendoza and Azizi, 2021; Peplowski and Marsh, 1997; Roberts, 2016; Roberts and Azizi, 2011; Roberts and Marsh, 2003; Rosario et al., 2016; Sawicki et al., 2015), but not in Borelli’s and Bennet-Clark’s law; indeed, as is discussed briefly later, in-series elasticity is usually framed as an evolutionary adaptation that overcomes the limits prescribed by Bennet-Clark’s law. And third, throughout this work, it is assumed that the contraction is inertial, i. e., that all opposing (parasitic) forces are small compared to the muscle force, and can thus be neglected. Parasitic forces are not considered in the classic presentation of either Borelli’s or Bennet-Clark’s law, and do not affect the conclusions presented here; a more general treatment may be found in Labonte (2023); Labonte et al. (2024); Polet and Labonte (2024). The three omissions have two important consequences: the first two imply that the speed of the payload is equal to the shortening speed of muscle at all times; and the third implies that the net work is equal to the work done by the muscle, and that all of this work flows into kinetic energy. Keeping both consequences in mind will help to follow the analysis, and is important when the results presented here are generalised, or compared to the more complex analyses presented elsewhere (Labonte, 2023; Polet and Labonte, 2024).

In discussing the limits to muscle mechanical performance, it can be physically insightful and biologically meaningful to explicitly assess the influence of animal body size. To facilitate such analyses, we will make the parsimonious assumption of isogeometry and isophysiology throughout this text. That is, in keeping with classic scaling theory, characteristic lengths, areas and volumes are assumed to scale with body mass *m* as *L* ∝ *m*^1*/*3^, *A* ∝ *m*^2*/*3^, and *V* ∝ *m*; and physiological parameters such as maximum stress, strain and strain rate are assumed to be size-invariant (Askew and Marsh, 2002; Close, 1972; Marden and Allen, 2002; Rospars and Meyer-Vernet, 2016).

## Results and Discussion

### Arbitrary speed with equal power — a problem of dimensions

Borelli’s and Bennet-Clark’s law both seek to estimate the maximum energy muscle can deliver with a single contraction. The key difference between them can be illustrated with a dimensional argument: if the left-hand side of an equation is of dimension length per time [L T^-1^], the right-hand side ought to be, too, otherwise an error has been made. Consider, then, the task of predicting the energy a muscle with fixed work and power capacity can deliver as it contracts against a mass *m*. Energy and work share the same dimension [M L^2^ T^-2^], so that, with Borelli, one may quickly surmise *E* = *W*_max_, or, in keeping with the focus of the jumping literature on take-off speed, 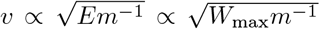. Note well that the only way to change *E* under a work constraint is to change *W*_max_—a variation of the work capacity is both necessary and sufficient to change the maximum possible energy output, and Borelli’s law thus encodes an energy-limit proper. Power, in contrast, is of dimension [M L^2^ T^-3^]—it is consequently impossible to write the energy output as a sole function of *P* (or define a speed using only *P* and *m*). Instead, dimensional consistency demands that one of three auxiliary variables be specified in addition: (i) a displacement *δ* of dimension [L], so that *E* ∝ (*P*_*max*_*δ*)^2*/*3^*m*^1*/*3^ (and *v* ∝ (*P*_*max*_*δm*^−1^)^1*/*3^); *or* (ii) a time *t* of dimension [T], so that *E* ∝ *Pt* (and 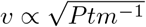); *or* (iii) a force *F* of dimension [M L T^-2^], so that *E* ∝ *P* ^2^*F* ^−2^*m* (and *v* ∝ *PF* ^−1^; see SI and Fig. 1 a-b). A crucial conclusion follows at once: a variation in power capacity is neither necessary nor sufficient to vary muscle energy output; changes in a third variable can alter the energy output resulting from the same power input, or keep it constant despite arbitrary variations in power input (Tab. 1).

**Table 1.**
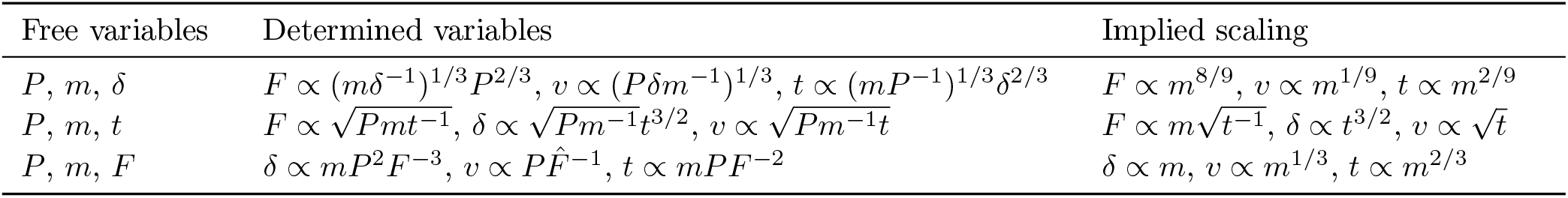
A dimensionally consistent link between power output *P*, payload mass *m*, and energy output *E* (or speed *v*) requires specification of one of three ‘auxiliary variables’: the contraction time, *t*, the contraction displacement, *δ*, or a characteristic force, *F*. Because point-mass dynamics involve five elemental variables, but provide only two independent equations, selecting any three variables fully determines the remaining two. The different instantiations of Bennet-Clark’s law differ in their choice of auxiliary variable, and consequentially in how the same mass-specific power output is partitioned into muscle force vs. muscle shortening speed—they thus yield different predictions for the allometry of muscle energy output and speed (see also Fig. 1 a-b). The choice of the auxiliary variable also controls the scaling of the remaining variables, as illustrated with predictions derived from isogeometry and isophysiology, i. e., *δ* ∝ *m*^1*/*3^, *F* ∝ *m*^2*/*3^, *P* ∝ *m*, and 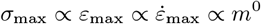.

**Figure 1.**
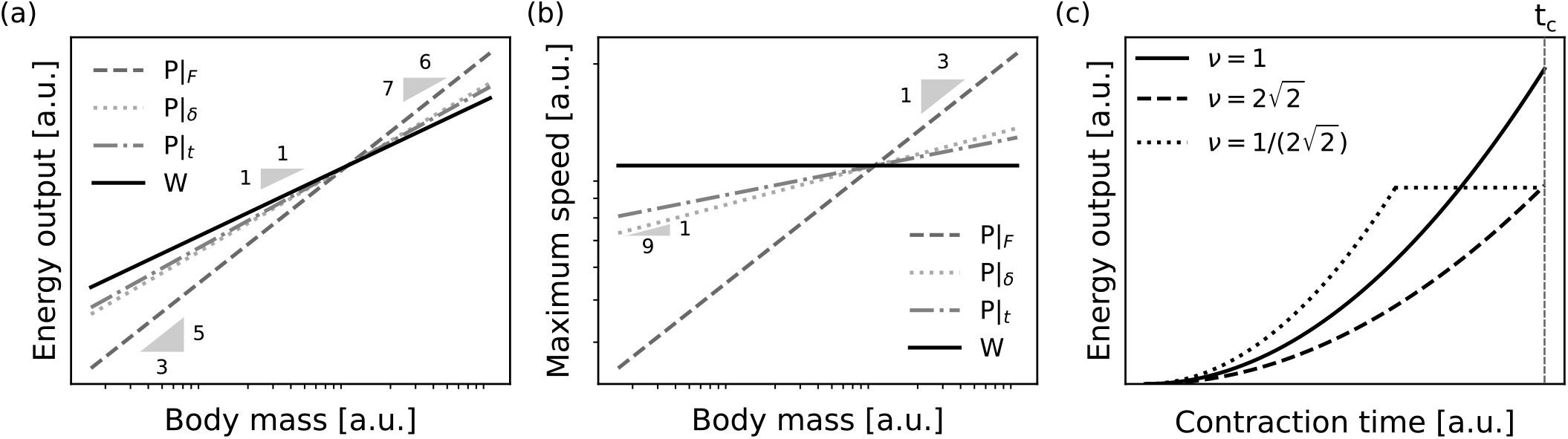
The assertion that muscle work and power capacity prescribe independent limits on muscle energy output has the corollary that changes in work or power capacity are necessary and sufficient to change muscle energy output. Using dimensional analysis, it can be confirmed that the muscle work capacity unambiguously defines a limit to (a) energy output and (b) payload speed (solid black lines). A dimensionally consistent link between energy, power capacity, payload mass and speed, however, requires specification of at least one additional auxiliary variable: a force *F*, a displacement *δ*, or a time *t*. Thus, a variation in muscle power capacity is neither necessary nor sufficient to achieve variations in muscle energy output or payload speed. Instead, different choices of auxiliary variable can lead to different, and indeed *arbitrarily different* predictions, as illustrated here by evaluating the energy output and speed for a muscle that delivers the same power capacity *P*, in combination with either an isogeometric and isophysiological force, *P*|_*F*_ (dark grey, dashed), displacement *P*|_*δ*_ (light grey, dotted), or time *P*|_*t*_ (medium grey, dash-dotted, see text for details). Borelli’s law thus encodes an energy limit proper, but Bennet-Clark’s law prescribes a combined power-force, power-displacement, or power-time constraint. (c) Even with both power capacity and auxiliary variable specified, the resulting energy output cannot be uniquely determined, as illustrated here with an example of three different muscles, all with the same volume and time-averaged power capacity, 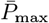, but with a different split into fascicle length, *l*_*m*_ and physiological cross-sectional area, *A*_*m*_; the muscles have a different aspect ratios, 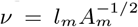. The same power is thus split differently into force and speed capacity. Let these muscles contract against a payload of mass *m*, for no more than a time *t*. How much energy can they inject? Because all muscles have the same power capacity, it is tempting to conclude that 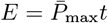, but all that can be said is 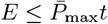 (see text; for simplicity, the plot illustrates a contraction for which the force is constant, but it can be generalised to any FVR). This limitation arises because Newtonian point mass dynamics only have three degrees of freedom; linking energy output, payload mass, time-averaged power and time thus also places demands on the muscle’s time-averaged force and shortening speed capacity (see text). Note that (a) and (b) are logarithmic, but (c) is on linear axes.

This simple observation has not-so-simple implications. A muscle’s work and power capacity depend solely on the muscle’s volume, and a characteristic stress, strain and strain rate. It thus seems reasonable to expect that specifying these quantities is all that is needed to predict the energy output with Borelli’s and Bennet-Clark’s law. For Borelli’s law, this is indeed so, but to estimate the energy output with Bennet-Clark’s law, the force and displacement capacity must be known, too—it becomes necessary to specify how a muscle volume *V* is split into physiological cross-sectional area, *A*_*m*_, and fascicle length, *l*_*m*_. In other words, the energy output now also depends on the muscle aspect ratio, 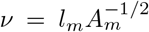 (Labonte, 2023; Polet and Labonte, 2024, This remains true if the time is fixed, see Fig. 1 c and below). Even with the muscle’s geometrical arrangement specified, the difficulties are not yet quite over—which of the three auxiliary variables should be chosen? The decision is not obvious, and lo!, examples for each option can be found: Bennet-Clark picked the displacement (Bennet-Clark, 1977), which remains the most popular implementation (e. g. Biewener and Patek, 2018; Bobbert, 2013; Gabriel, 1984; James et al., 2007; Marsh, 1994; Sutton et al., 2016). Usherwood instead fixed the time (Usherwood, 2013; Usherwood and Gladman, 2020), and, last but not least, Meyer-Vernet and Rospars (Meyer-Vernet and Rospars, 2015, 2016) and Hawkes et al (Hawkes et al., 2022) fixed the force. The specific choice carries meaningful consequences: it leads to quantitative differences in the downstream performance prediction. Isogeometry and isophysiology imply *δ* ∝ *m*^1*/*3^ and *F* ∝ *m*^2*/*3^, which leads to *E* ∝ *m*^11*/*9^ or *E* ∝ *m*^5*/*3^, and *v* ∝ *m*^1*/*9^ or *v* ∝ *m*^1*/*3^, respectively (Bennet-Clark, 1977; Meyer-Vernet and Rospars, 2015). Usherwood instead assumed 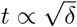, leading to *E* ∝ *m*^7*/*6^ and *v* ∝ *m*^1*/*12^ (Usherwood, 2013; Usherwood and Gladman, 2020, or, via a similar argument, *E* ∝ *m*^4*/*3^, and *v* ∝ *m*^1*/*6^). A rather striking difference between the two laws has become apparent: Borelli’s-law unequivocally predicts *E* ∝ *m* and *v* ∝ *m*^0^ = constant. Bennet-Clark’s law, however, has been used to predict anything between *E* ∝ *m*^7*/*6^ to *E* ∝ *m*^5*/*3^, and *v* ∝ *m*^1*/*12^ to *v* ∝ *m*^1*/*3^ (Table 1 and Fig. 1 a-b). In fact, it can predict *any* energy output through suitable variation of the auxiliary variable of choice (Fig. 1 a-b). It is evidently not the muscle’s power capacity itself that is limiting the energy output, and Bennet-Clark’s *laws* may thus at best be said to encode a power-displacement, power-time, or power-force constraint. However, and no less clearly, the energy output of muscle can, in fact, violate Borelli’s law. If it is neither the muscle’s work nor power capacity that is imposing the limit in these instances, then what is?

### Power-limits and the hidden determination of centre-of-mass dynamics

In the previous section, it was demonstrated that a dimensionally consistent link between a fixed power input and energy output requires specification of one auxiliary variable, that this choice is not obvious, and that different choices lead to different results. The task that lies ahead is to identify the mechanical explanation for these differences.

Consider an animal of body mass *m*. Let the maximum time-averaged power capacity of its muscles be 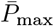, and allow it to accelerate for no more than a time *t*. What is the energy imparted to its centre-of-mass? This is a classic textbook setup for Bennet-Clark’s law, and it is tempting to conclude 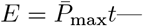but this is not necessarily so. In fact, all that can be deduced is the considerably weaker 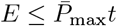 (Fig. 1 c). This restriction arises because specifying the body mass *m* together with 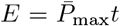 places hidden demands on the muscle force and shortening speed capacity: it requires that the time-averaged force capacity is at least 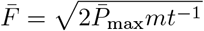, and that the maximum shortening velocity is at least 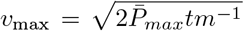. If the force capacity is smaller, the muscle cannot deliver the power 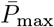 within time *t*; and if its shortening speed capacity is smaller, it can deliver *P*_max_, but in less time. In both cases, 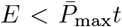 (Fig. 1 c). In other words, although the maximum average power capacity is equal to 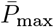 no matter the muscle aspect ratio, only one unique aspect ratios allows a muscle with a maximum 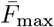 and *v*_max_ to deliver 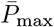 exactly within time *t* (Fig. 1 c). This point may appear subtle, but its consequences are surely troubling: estimating the energy output associated with a specific muscle power input requires to specify an auxiliary variable by physical necessity—but even that may still not yield a definite answer.

To understand the mathematical origin of this result, note that Newtonian point-mass dynamics are governed by a set of two equations that uniquely link five elemental variables: Newton’s second law and its path-integral define the relation between force, mass, speed, time, and displacement throughout the contraction. This mathematical structure dictates that the choice of any three parameters uniquely determines the remaining two variables. Thus, regardless of how Bennet-Clark’s law is implemented for an animal with body mass *m, all* dynamic variables end up fully defined (Tab. 1). Contrast this scenario with a determination of the energy output via the work capacity of muscle, which requires to define only one additional parameter, *E* = *W*_max_. It consequently does not matter whether *W*_max_ is partitioned into a small force and large displacement capacity (which would take a long time, and involves low power), or into a large force and a small displacement capacity (which will be completed rapidly, and requires large power)—any muscle with work capacity *W*_max_ will do, because the governing equations remain underdetermined, and retain one residual degree of freedom that can absorb arbitrary work partitioning. It may be tempting to file this observation as technically correct, but of limited practical implication. This would be a mistake. Consider again the most widespread quantitative implementation of Bennet-Clark’s law, which combines a size-invariant power density with an isogeometric displacement to predict *E* ∝ *m*^11*/*9^ and *v* ∝ *m*^1*/*9^. Hidden within this prediction lies the necessary condition that the average muscle force scales as *F* ∝ *m*^8*/*9^, in substantial excess of the isogeometric and isophysiological expectation, *F* ∝ *m*^6*/*9^ (Tab. 1). How this positive allometry may be achieved in an isogeometric and isophysiological system is not obvious.

Dimensional arguments and the mathematics of point mass dynamics in combination provide two conclusions: due to the need for dimensional consistency, the question “how much energy can a motor with power *P*_max_ inject into a mass *m*” cannot be answered without specification of exactly one further auxiliary variable (Tab. 1); and due to the fundamental structure of Newtonian dynamics, any such choice uniquely defines all remaining variables—the muscle force, displacement and shortening speed throughout the contraction are fully determined. It is the determination of the shortening speed in particular that is the essential distinction between the different instantiations of Bennet-Clark’s law, and that brings about the variation in the energy output they predict—an assertion to which the discussion will now turn.

### Beyond power-limits: The kinetic energy capacity of muscle

The notion that muscle power capacity limits muscle energy output has been called into question in the past (e. g. Adamson and Whitney, 1971; Farley, 1997; Knudson, 2009; Ruddock and Winter, 2015; Winter, 2005; Winter et al., 2016). But the harshness with which this criticism was sometimes expressed masked its own failure to address the fundamental issue and valid concern unearthed by the careful observations of Bennet-Clark and many others since: any account of muscle mechanical performance that ignores the dimension of time risks to arrive at conclusions that violate physiological or physical constraints, for no muscle can do a unit of work in arbitrarily short contraction times. It is true enough that the mechanical quantity that uniquely ties speed, mass, and contraction time is the impulse, and not power (Adamson and Whitney, 1971; Knudson, 2009; Ruddock and Winter, 2015; Winter, 2005; Winter et al., 2016); but pointing this out merely addresses a symptom instead of the problem’s root: what limits the time over which muscle can do work?

Because energy is the focal metric in both Borelli’s and Bennet-Clark’s law, it is convenient as much as reasonable to approach this question via the conservation of energy— the path-integral of Newton’s second law:

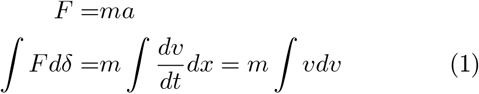

This writing reveals immediately and unambiguously that there exists a scenario for which the energy output is not determined by the muscle’s work capacity: when the muscle reaches its maximum contraction speed before it has exhausted its displacement capacity (Fig. 2 Labonte, 2023, In the SI, the same result is derived via the time-integral, i. e., through explicit consideration of the muscle’s *impulse capacity*). To unpack this assertion, note that evaluation of eq. 1 requires specification of one of two integration boundaries: a maximum displacement, *δ*_max_, *or* a maximum speed, *v*_max_ (Labonte, 2023). Fixing the displacement is the usual choice; the energy output is then limited by the muscle’s work capacity, and one concludes with Borelli that *E* = *W*_max_ ∝ *m*. However, muscle has not only a maximum shortening distance, but also a maximum shortening speed (Hill, 1938)—it is thus no less logical to fix the upper bound for velocity integral, which yields the “Hill-limit”, 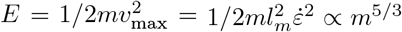 (Labonte, 2023; Labonte et al., 2024). The Borelli- and the Hill-limit can clearly differ, and they thus represent two independent constraints on the muscle’s ability to do mechanical work. Muscle has not one, but *two* characteristic energy capacities—the work capacity, 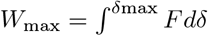 is joined by the no less fundamental *kinetic energy capacity*, 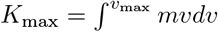 (Labonte, 2023)

**Figure 2.**
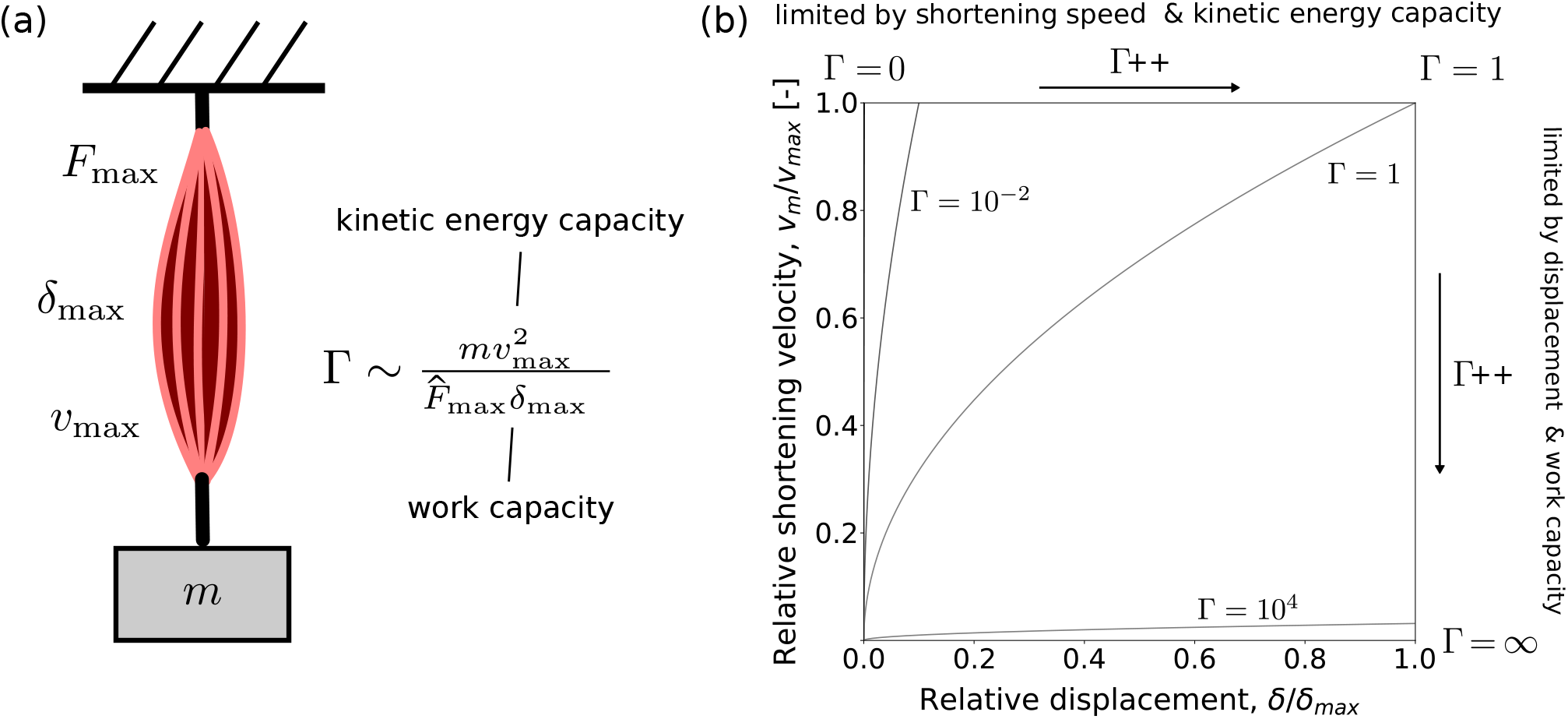
**(a)** An idealised musculoskeletal system is characterised by a maximum muscle force *F*_max_, a maximum muscle displacement capacity, *δ*_max_, and a maximum muscle speed of shortening, *v*_max_. Combined with the payload mass *m*, this mechanical system has four dimensional parameters, but point mass dynamics only permit to specify three. Which parameters are free and which are fixed is determined by the magnitude of the dimensionless physiological similarity index, Γ. **(b)** One interpretation of Γ emerges from the inspection of “equation of motion landscapes”, in which the dynamic progression of the displacement and speed is monitored as the muscle contracts (Labonte, 2023). For Γ → 0, muscle acquires shortening velocity rapidly and with a minimal fraction of its displacement capacity. The contraction always ends with maximum shortening speed, but involves variable muscle displacement; it becomes quasi-instantaneous. For Γ → ∞, the muscle has contracted by its maximum displacement long before it has reached any appreciable fraction of its maximum shortening speed. The muscle always shortens maximally, but achieves variable shortening speeds; the contractions becomes quasi-static. The transition from a shortening speed to a displacement limit occurs at a limiting value of Γ = 1, the critical value at which muscle reaches the maximum displacement and shortening speed at exactly the same time. The EoM landscape here is for a muscle that has a force-velocity relationship (FVR) idealised as a step-function; a generalisation of the concept to any FVR can be found in Labonte (2023).

*W*_max_ and *K*_max_ could be equal, but there is no fundamental reason why they would have to be—they depend on different physiological processes and mechanical properties. The limit to muscle energy output is thus, in general, set by whichever of the two characteristic energy capacities is smaller. To identify the relevant limit, it is convenient to evaluate their ratio, Γ (Fig. 2; Labonte, 2023):

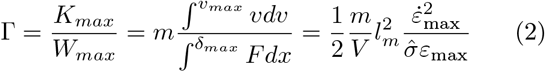

For Γ ≤ 1, the kinetic energy capacity is limiting, and for Γ ≥ 1, the work capacity is limiting (Fig. 3). So how large is Γ? This question is of obvious and immediate importance, and the subject of the recently developed theory of physiological similarity (Labonte, 2023; Labonte et al., 2024; Polet and Labonte, 2024), from which the following analysis draws.

**Figure 3.**
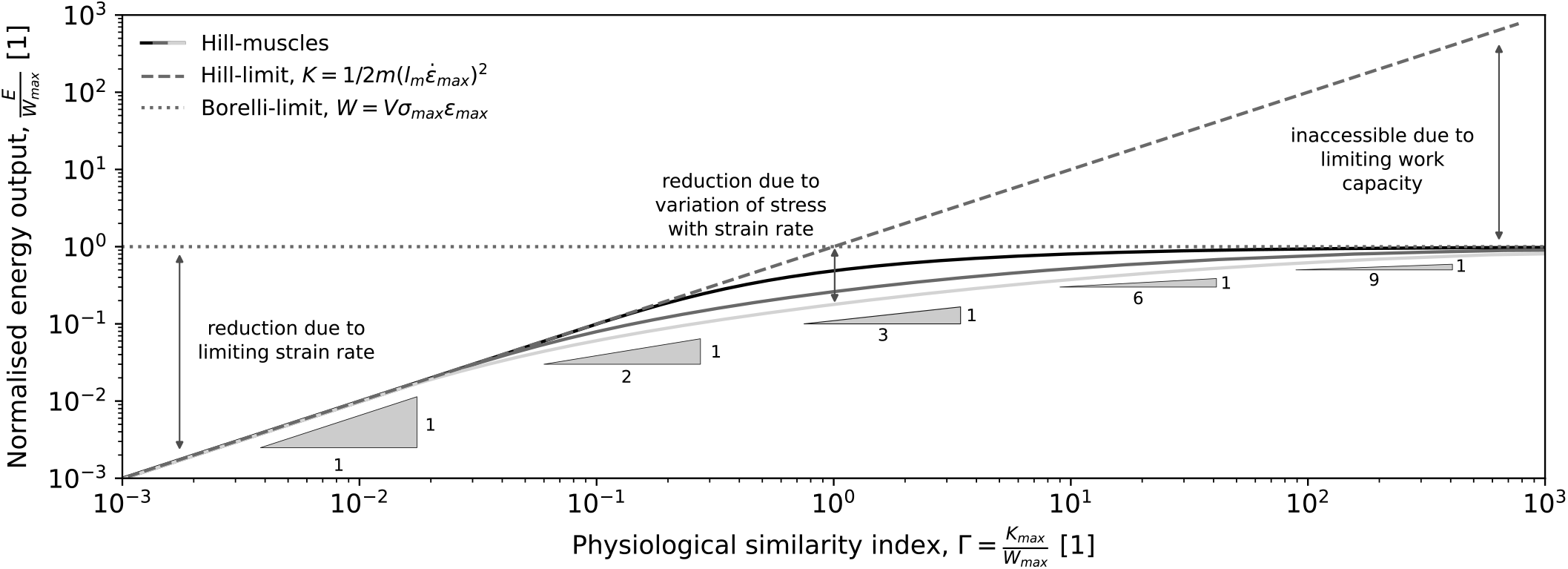
Muscle is characterised not by one, but by two characteristic energy capacities: the work capacity, *W*_max_, established by Borelli, is joined by the no less fundamental kinetic energy capacity, *K*_max_. Because the kinetic energy capacity stems from a limit on the maximum muscle strain rate, it is also referred to as the Hill-limit to muscle energy output in this text (Labonte, 2023). The ratio of both energy capacities defines the dimensionless physiological similarity index, Γ = *K*_max_*/W*_max_, which is a suitable proxy for the mechanical energy muscle can deliver. For Γ *<<* 1, *E* ≈ *K*_max_ and the muscle’s kinetic energy capacity is limiting; and for Γ *>>* 1 the muscle is limited by its work capacity, *E* ≈ *W*_max_. The ability of muscle to deliver energy is further reduced by the variation of stress with strain rate, as described via the Hill-relation (eq. 4). The three solid lines show the results for *Q* = 0 (a linear force-velocity-relationship, shown in black), *Q* = 4 (a typical value for vertebrate muscle; dark grey Alexander, 2003), and *Q* = 10 (an extreme value, lightgrey). The effect of a Hill-type force-velocity-relationship (FVR) on energy output is small compared to the constraint imposed by the kinetic energy and maximum work capacity for sufficiently small and sufficiently large Γ, and is maximal for Γ = 1. In many cases, an estimation of muscle energy output via the analytically simple Hill- and Borelli-limits will thus provide a robust first order estimate. However, the effect of Hill-type FVR is important, too: it results in a more complex relationship between energy output and Γ as indicated by the slope triangles (eq. 6).

Equation 2 is completely general in the sense that it holds for any muscle that is restricted by a maximum stress, strain, and strain rate, i. e. regardless of the exact shape of the force-velocity relationship (FVR; Labonte, 2023; Mendoza et al., 2023). But the reader will rightfully point out that this generality does little good, for eq. 2 depends not on *σ*_max_, but strictly on 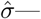the muscle stress *averaged over the exerted strain*—and the exact relationship between these two quantities does necessarily depend on the specific form of the FVR. FVRs thus influence Γ in two distinct ways: through the imposition of a maximum strain rate, encoded via 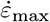 in the numerator; and through the variation of stress with strain rate, implicit in the appearance of 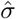 in the denominator—in other words, they influence both *K*_max_ and *W*_max_. What is the relative importance of these two FVR features in determining muscle energy output?

To evaluate the effect of a maximum strain rate independent of the effect of a variation of stress with strain rate, the FVR may be idealised as a step function; the muscle stress is independent of the strain rate until the maximum strain rate is exceeded, at which point it drops instantaneously to zero (Ilton et al., 2018; Labonte, 2023; Polet and Labonte, 2024). For such an idealised muscle, 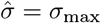, and thus:

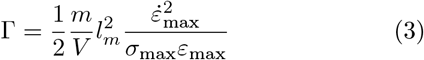

To quantify the additional effect of the variation of muscle stress with strain rates below 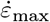, the FVR may be described instead via a normalised Hill-relation (Hill, 1938; McMahon, 1984):

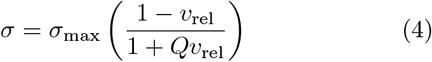

where *Q* is a dimensionless constant, typically of order unity, and 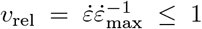 is the relative strain rate. Finding an explicit symbolic expression for the displacement-averaged stress such a muscle can generate is, to the best of our judgement, only possible for *Q* = 0, i. e., a linear FVR, for which Γ reads (see SI):

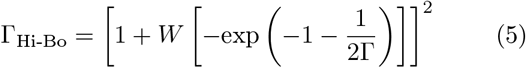

where W is the productlog- or LambertW-function, and where the subscript indicates the Hill-Borelli-limit (Labonte et al., 2024).

Comparison of eqs. 3 and 5 provides the conclusion that the energy output of both the idealised and the Hill-type muscle is bound by the kinetic energy capacity, 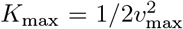 for sufficiently small, and by the maximum work capacity, *W*_max_ = *V σ*_max_*ε* for sufficiently large Γ (Labonte, 2023, for a more detailed discussion of the differences, see SI). Using numerics, it can be validated that this observation generalises for any value of *Q* (Fig 3); how exactly stress varies with strain rate is irrelevant for both very small and very large Γ, because contractions become quasi-instantantenous or quasi-static, respectively (Fig. 2; Labonte, 2023).

To evaluate the energy output at intermediate values of Γ, note that for the idealised muscle, *E* = *K*_max_ if Γ ≤ 1, and *E* = *W*_max_ for Γ ≥ 1 (Labonte, 2023). For a Hill-type FVR with *Q* = 0, however (see SI):

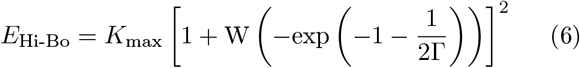

The two predictions are practically identical for both small and large Γ; they differ by no more than a factor of about two at Γ = 1 (for *Q* = 4, a typical value for vertebrate muscle, this difference increases to a factor of about four, and for an extreme value of *Q* = 10, it is a factor of about six, Fig. 3). This result has both physical and practical implications: the magnitude of Γ says something meaningful about the ability of muscle to deliver mechanical energy regardless of the exact form of the FVR; the limit to energy output in small animals arises primarily from the existence of maximum muscle strain rate, and not from the variation of stress with strain rate as encoded by the Hill-relation (Bobbert, 2013; Sutton et al., 2019); and the muscle energy can often be evaluated with reasonable accuracy through the simple expressions that define the Hill- and the Borelli-limit for an idealised FVR. All this is not to say that the variation of stress with strain-rate does not have meaningful implications—some examples are illustrated further below.

With these results at hand, it is now finally the time to discuss the magnitude Γ as defined by eq. 3, and to thus answer the question which of the two characteristic muscle energy capacities, the Borelli- or the Hill-limit, may be relevant in animal movement. Several case studies can be found in the literature (Labonte, 2023; Labonte et al., 2024; Polet and Labonte, 2024). As an illustrative example, consider the musculoskeletal system that propels running animals, for which Γ ≈ 0.07 mass^2/3^ kg^-2/3^ (Labonte et al., 2024). Thus, from a 0.1 mg mite to a 10 t elephant, Γ is predicted to vary by a whopping seven orders of magnitude (Labonte, 2023; Labonte et al., 2024). It remains smaller than unity for runners lighter than about 50 kg (Labonte et al., 2024), and the kinetic energy capacity, *K*_max_, is thus likely a robust proxy for the limit to muscle energy output in the vast majority of terrestrial animals (Labonte, 2023; Labonte et al., 2024).

### Biological relevance and testable predictions

Framing the limits on muscle energy output in the conceptual terms of a limiting work vs. power density has become a textbook staple, invoked to explain an extraordinarily diverse array of observations in comparative animal biomechanics. Our analysis reveals that attributing these observations to a competition between a limiting work and power density is to miss out on some important physics, and that the alternative constraint on muscle energy output is not imposed by the muscle’s power density, but by its characteristic kinetic energy density, *K*_*ρ*_ = *K*_max_*/m*_*m*_. What are the implications of this conclusion for our understanding of the biomechanics of animal movement?

*K*_*ρ*_ differs from the power density *P*_*ρ*_ and the work density *W*_*ρ*_ in at least four aspects, and these differences provide clear and consistent explanations for some classic observations in comparative animal biomechanics. First, in contrast to *P*_*ρ*_ and *W*_*ρ*_, *K*_*ρ*_ retains a size-dependence (Figure 4 a): geometrically and physiologically similar larger musculoskeletal systems have a larger kinetic energy density, *E*_*ρ*_ ∝ *m*^2*/*3^. As a result, larger animals are generally faster (Bejan and Marden, 2006; Garland, 1983; Gazzola et al., 2014; Labonte et al., 2024; Meyer-Vernet and Rospars, 2016; Pennycuick, 1968; Sánchez-Rodríguez et al., 2023). Second, in contrast to *P*_*ρ*_ and *W*_*ρ*_, *K*_*ρ*_ can be geared, 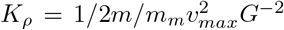, where *G* is a dimensionless mechanical advantage, defined as the ratio between system output and muscle input force (Figure 4 b, Labonte, 2023; Polet and Labonte, 2024). As a result, two staples of biomechanical analyses that may ring contradictory can in fact both hold true: gearing is usually interpreted in terms of force-velocity trade-offs; a lower gear ratio increases the instantaneous velocity of the payload at the expense of the transmitted force. But gearing leaves the work and power capacity unaffected, for it amplifies displacement and velocity by just as much as it attenuates force. How, then, can a muscle make things move more quickly via gearing, although its putatively limiting work and power capacities have remained unchanged? The answer is that for as long the energy output remains below the work capacity, a reduction of the gear ratio can in fact enable muscle to do more work, because it increases its kinetic energy capacity (McHenry, 2011, 2012; Olberding et al., 2019; Osgood et al., 2021; Polet and Labonte, 2024). The immediate implication of this observation is the existence of a mechanically optimal mechanical advantage that varies with animal size and environment—a hypothesis unpacked in detail in Polet and Labonte (2024). Third, in contrast to *P*_*ρ*_ and *W*_*ρ*_, *K*_*ρ*_ is a function of the mass that is driven (Figure 4 c). This is perhaps the least intuitive idiosyncrasy of *K*_*ρ*_: *increasing the payload can enable muscle to deliver more energy*, because it unleashes latent work capacity (Fig. 3, and see Sawicki et al., 2015, for a related finding on muscle power output). As a result, where *K*_*ρ*_ is limiting, animals may be able to achieve the same speed for payloads that are increasing multiples of their own body mass. As a striking illustration of this prediction, consider rhinoceros beetles, which can carry up to 30 times their own body mass without changing speed (Kram, 1996). Fourth, the alleged power-limit to muscle energy output is often invoked to explain a key functional benefit of in-series elasticity in musculoskeletal systems: in dynamic contractions, tendons can decouple limb and muscle shortening speed, and muscle can consequently achieve similar absolute limb speeds with lower muscle shortening speeds, so increasing its power output (Aerts, 1997; Astley and Roberts, 2012; Farris et al., 2016; Galantis and Woledge, 2003; Kurokawa et al., 2001; Marsh, 2022; Roberts and Marsh, 2003; Robertson et al., 2018); in quasi-static “latched” contractions, muscle can contract arbitrarily slowly against elastic elements, and so avoid both force-velocity-effects and supposed muscle power limits to performance, by instead releasing its work capacity explosively (Bennet-Clark, 1975; Bennet-Clark and Lucey, 1967; Gronenberg, 1996; Longo et al., 2019; Patek, 2023). A large body of careful work has been dedicated to such amplification of muscle power, be it in dynamic or in “latched” quasi-static contractions (for recent reviews, see Holt and Mayfield, 2023; Longo et al., 2019; Patek, 2023). There is no doubt, of course, that elastic elements can amplify muscle power. But a reasonable argument is to be had whether the biological function of these “springs” in these instances is to amplify speed rather than power as such. The kinetic energy capacity of a spring is likely orders of magnitudes higher than that of muscle; it is limited by the elastic wave speed, 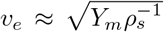, where *Y*_*m*_ is the Young’s modulus of the spring, and *ρ*_*s*_ is its density. For reasonable values of *Y*_*m*_ ≈ 10^9^*Nm*^−2^ and *ρ*_*s*_ ≈ 1000*kgm*^−3^, one finds *v*_*e*_ ≈ 1000 m s^-1^; a muscle with a typical maximum strain rate of 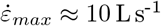 would need to have fascicles with a length of 100 m to reach the same absolute speed. Because jumping performance in small animals is likely limited by the kinetic energy capacity of muscle, we posit that (i) their springs act as “work enablers” (see also Roberts and Marsh, 2003), allowing them to overcome the constraint on energy output imposed by a limiting kinetic energy capacity; and (ii) that power amplification is an epiphenomenon instead of the biological purpose of in-series elasticity in rapid movements. The outcome of this somewhat semantic debate is clearly immaterial for the validity of the long list of fundamental insights that have been derived from the study of power amplification due to biological “springs” (Gronenberg, 1996; Ilton et al., 2018; Longo et al., 2019; Patek, 2023).

**Figure 4.**
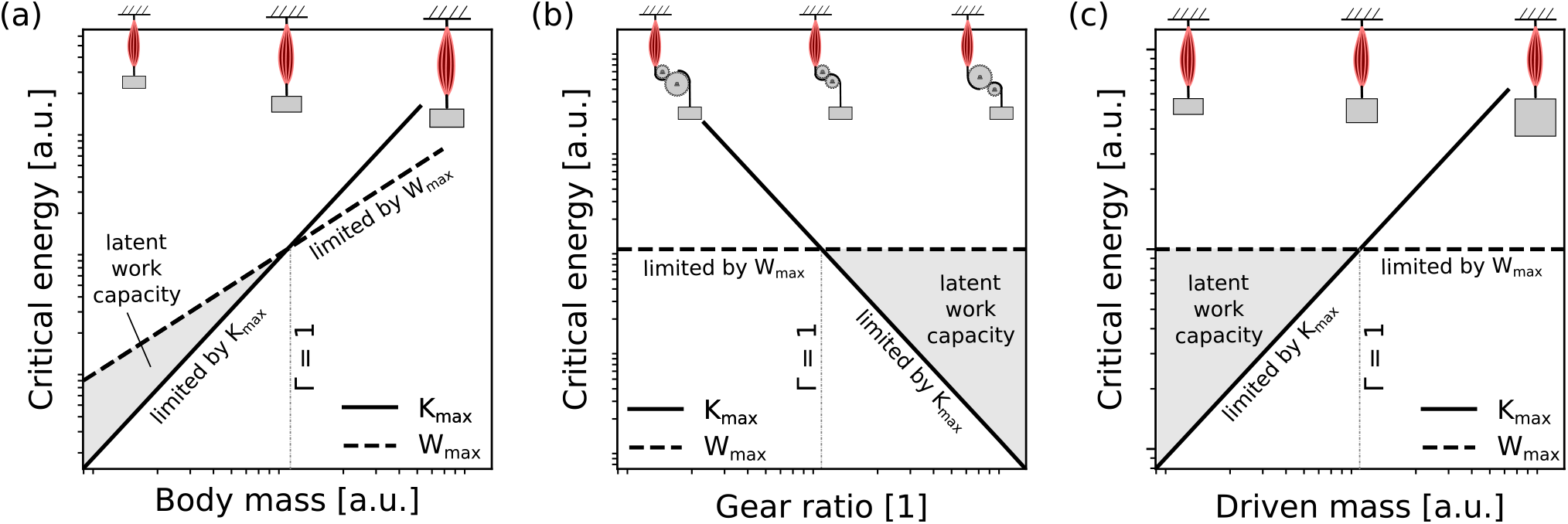
The energy output of muscle is limited by two critical energy capacities: the maximum *work capacity, W*_max_ = *V σ*_max_*ε*_max_, and the *kinetic energy capacity*, 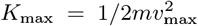. Three differences between *K*_max_ and *W*_max_ have noteworthy implications for the variation of biomechanical performance across animal size and musculoskeletal “design”. **(a)** Both *K*_*max*_ and *W*_*max*_ increase with size for geometrically similar musculoskeletal systems, but at different rates, *W*_*max*_ ∝ *m* vs. *K*_*max*_ ∝ *m*^5*/*3^. As a consequence, small animals are more likely to be limited by their kinetic energy capacity, and large animals are generally faster (Garland, 1983; Labonte et al., 2024). **(b)** The work capacity is unaffected by changes to the gear ratio *G, W*_*max*_ ∝ *G*^0^, but gearing changes the kinetic energy capacity, *K*_max_ ∝ *G*^−2^. As a consequence, where the energy output is limited by the kinetic energy capacity, it can be increased by changing the gear ratio, such that small animals benefit from smaller gear rations, and large animals benefit from larger gear ratios (Labonte, 2023; Polet and Labonte, 2024). **(c)** For the same musculoskeletal system, the work capacity of muscle is independent of the payload, *W*_*max*_ ∝ *m*^0^, but the kinetic energy capacity is directly proportional to it, *K*_max_ ∝ *m*. As a consequence, animals that are limited by *K*_max_ can respond to an increase in payload by delivering more energy; the increase in load releases latent work capacity. All three characteristics of *K*_max_—its dependence on animals size, gear ratio, and payload—distinguish it meaningfully from the power capacity of muscle, and so sharpen the physical explanation of several observations in comparative movement biomechanics (see text).

The above examples may perhaps sharpen the physical explanation of some well-established observations in the comparative biomechanics of animal movement, but they do not make novel performance predictions as such. To derive such predictions, we next compare the theory of physiological similarity directly to classic scaling theory.

The importance of animal size in determining physiology, morphology and physical constraints is well-established, and perhaps among the oldest and most intensely studied aspects of comparative biomechanics (McMahon et al., 1983; Schmidt-Nielsen, 1984, For a recent review, see Clemente and Dick (2023).). Where such inquiries are concerned with dynamics, they typically invoke a characteristic muscle force capacity, *F* ∝ *m*^2*/*3^, a characteristic displacement capacity, *δ* ∝ *m*^1*/*3^, and a characteristic work and power capacity, *W* ∝ *m* and *P* ∝ *m*, respectively. Together with the payload, classic scaling theory thus specifies four mechanical quantities— *m, σ*_max_, *ε*_max_, and 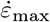 (Figure 2 a)—but point mass dynamics only provides *three* degrees of freedom. The startling consequence of this over-determination is that muscle is characterised not by one force, energy, speed displacement, and power capacity, as classic scaling theory would have it, but by *two* (Labonte, 2023). A full analysis of this observation exceeds the scope of this work, and will have to await further study; but it will be illustrated with one brief example.

In the preceding text, Γ was defined as the ratio of two characteristic energies. It can however be derived just as well as the ratio of two characteristic forces (Labonte, 2023; Polet and Labonte, 2024):

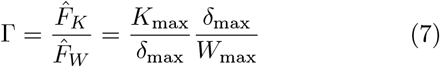

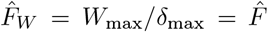 is the displacement-averaged muscle force associated with the muscle’s work capacity, and the typical attendant in classic scaling theory; 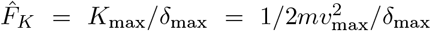, in turn, is the displacement-averaged inertial force associated with the muscle’s kinetic energy capacity, which has, to the best of our knowledge, escaped general attention. Importantly, the two characteristic forces differ in their dependence on animal body size: for an idealised muscle, 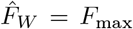, and, for Γ *>* 1, one recovers the classic isometric prediction, 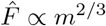. For Γ *<* 1, in turn, one finds 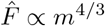— in substantial excess of textbook isometry. In practice— that is for a muscle with a Hill-type FVR—the realised scaling will fall between these two extremes (Fig. 5 b), as can be confirmed by inspecting the displacement-average force (see SI):

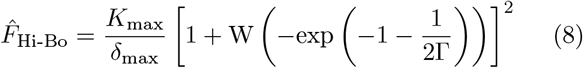

**Figure 5.**
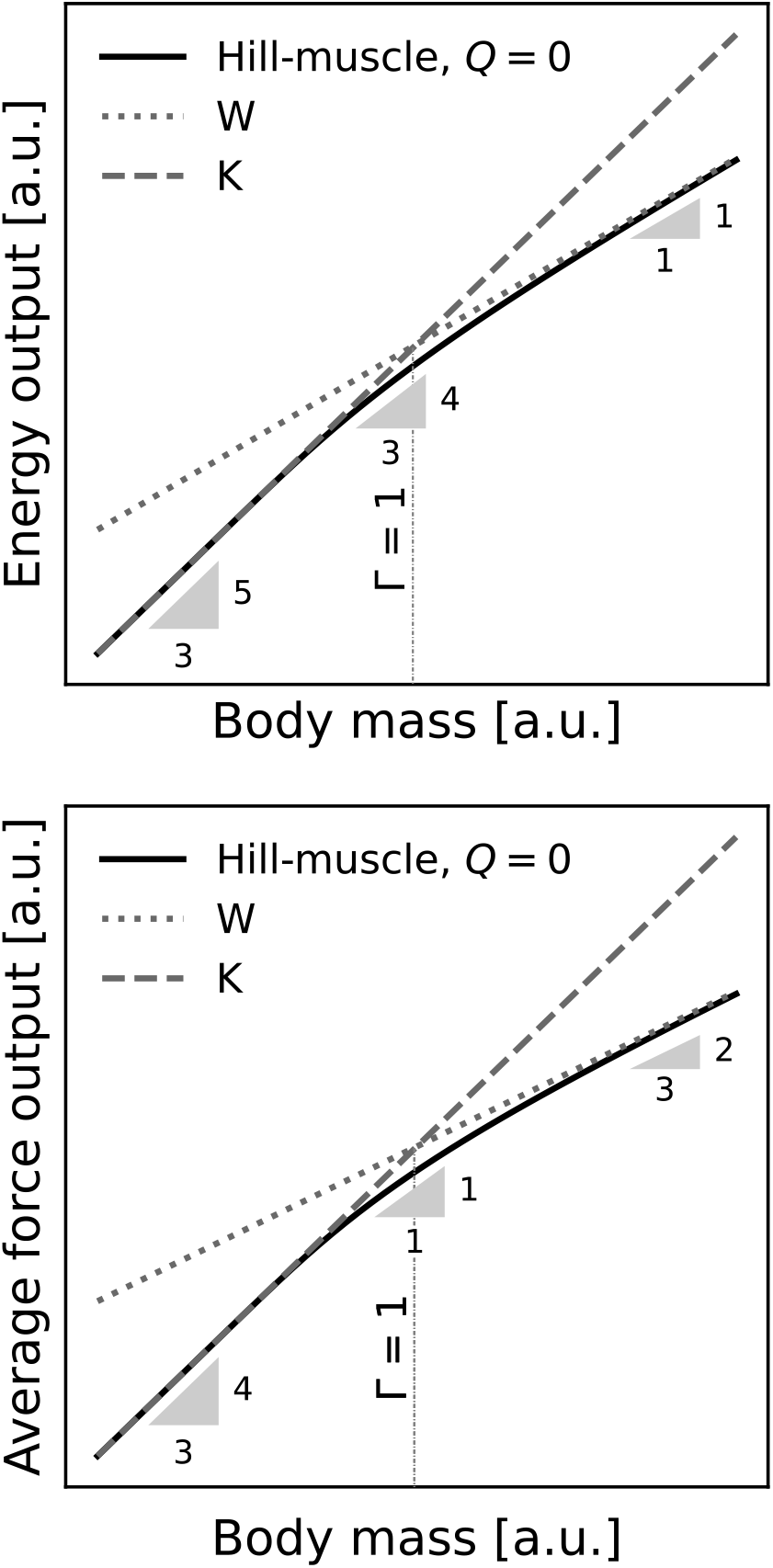
The theory of physiological similarity predicts that muscle has not one but two characteristic energy, force, speed, displacement and power capacities. This result arises because a typical musculoskeletal system is characterised by four dimensional quantities—a payload mass, and a force, displacement, and shortening speed capacity—but Newtonian dynamics only provides three degrees of freedom. The magnitude of Γ ∝ *m*^2*/*3^ does thus not only dictate the variation of (a) muscle energy output, but also of (b) the displacement-averaged force output with animal body mass *m*. Neither scaling relationship can be characterised satisfactorily by a single scaling coefficient— a notable difference to classic scaling theory. One consequence of this difference it that the maximum displacement-averaged force can grow with positive allometry even for isogeometric and isophysiological animals. Larger animals are thus able to do more mass-specific work, and thus move with larger absolute speeds (Labonte et al., 2024).

A remarkable conclusion seems inescapable: the predicted variation of the displacement-averaged force 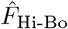 with animal size violates the prediction from classic scaling theory for all but perhaps the very largest animals (Fig. 5 b). This non-trivial scaling provides a hypothetical answer to a major outstanding question in the allometry of animal locomotor performance (Alexander, 2005): larger animals are generally faster, which implies that their muscles do more mass-specific work. Do they do so with a positively allometric force, a positively allometric displacement, or both? And how is this positive allometry achieved, given that musculoskeletal systems that vary substantially in size generally tend to conform to isogeometry and isophysiology? The answer that emerges from the theory of physiological similarity is that the muscle force averaged across an isometric displacement can grow with positive allometry even for isogeometric and isophysiological animals, because larger animals accelerate more slowly, and their muscles thus spend more time in favourable regions of the Hill-relation, where muscle force capacity is high (Figs. 2 and 5).

Equation 8 illustrates that the relevance of Γ is not restricted to energy output. Indeed, muscles that operate with equal Γ can be shown to deliver the same fraction of their maximum work and power capacity; to operate at the same fraction of their speed and displacement capacity; and to generate the same ratio of a characteristic inertial to a characteristic maximum force (Labonte, 2023; Labonte et al., 2024; Polet and Labonte, 2024). Because these parameters characterise a wide array of muscle physiological and mechanical characteristics—and to avoid giving one interpretation priority over any other— they may be emphasised equally by defining Γ as an index of *physiological similarity* (Labonte, 2023).

### A useful shift in perspective?

The idea that muscle power limits muscle energy output has become common biomechanical vernacular. It is rooted in the fundamental objection that muscle needs time to inject energy, and that muscle work capacity alone therefore does not tell the whole performance story. Supported by a series of arguments, we have suggested that this conclusion is partially wrong, and thus only partially right. Variations in muscle power are neither necessary nor sufficient to vary the energy muscle can deliver in a single contraction—a muscle’s power capacity consequently does not pose a limit to either energy output or speed by itself. The problem is not that small animals do not have enough time to do work, the problem is that their muscles have a lower maximum shortening speed, and consequently a lower kinetic energy capacity: neither infinite power nor an infinitely long contraction time would help overcome this limit. Power, of course, is not relegated to biomechanical irrelevance altogether; it is solely the assertion that muscle power capacity limits muscle energy output that is called into question.

A limit to shortening speed is tacit in any dimensionally consistent expression that links muscle power, mass and energy, and consequently shares many of the features that are typically associated with a power-limit, including shorter contraction time scales, and a reduced mechanical performance in small animals. The conclusion therefore neither can nor should be that the large body of work that analysed problems in biomechanics in the conceptual terms of a power-limit is “wrong” in its conclusions, that it lost any significance, or that the fundamental issues it raised are any less seminal. Much rather, the question ought to be whether a shift in perspective, to muscle shortening speed instead of muscle power, and to a kinetic energy density instead of power density, brings any meaningful advantages, or if it is at best technically correct, but for all intents and purposes practically irrelevant.

In favour of this shift, four brief arguments may be presented. First, expressing putative energy-limits directly in terms of energy enables meaningful comparison, for distinct limits now share the same dimension (Figure 4). Second, this comparison provides straightforward explanations for a series of observations in comparative biomechanics that are cumbersome if not impossible to explain in the framework of a size-invariant work and power density, and provides predictions for the scaling of musculoskeletal performance that depart from textbook theory (Figs. 4 and 5). Third, in its reliance on auxiliary variables, analysing muscle contractions in terms of muscle power makes it exceedingly easy to unintentionally demand of muscle something it may not be able to do. An explicit account of the key mechanical variables that limit every contraction — via the physiological similarity index, Γ — side-steps this difficulty, and provides a clear framework to ensure that mechanical analyses remain not only physically but also physiologically plausible. It is both clearer and less ambiguous to bind speed through an explicit limit on shortening velocity, than to introduce this limit through the backdoor, by treating the problem as if it were one of muscle power. Fourth, the notion of a size-invariant work and power density leaves remarkably little room for adaptive variation in musculoskeletal design. Inspection of the kinetic energy density, in turn, permits speculation. To give but two examples: (i) systematic variations in gear ratio with size can, in fact, enhance the work output of musculoskeletal systems, such that small animals would benefit from small, and large animals from large gear ratios (Biewener, 1989; Labonte, 2023; Labonte et al., 2024; Polet and Labonte, 2024; Usherwood, 2013); (ii) the maximum energy output can be independent of muscle mass, and instead depend solely on fascicle length, gear ratio, and the maximum muscle strain rate. Thus, small animals may be able to reduce the fraction of the body mass allocated to muscle, without suffering from a decrease in locomotor speed, as appears to be the case in reptiles compared to mammals (Labonte et al., 2024). These hypotheses no doubt require scrutiny, but they follow readily from inspection of the kinetic energy density, and cannot be easily extracted through the lens of a limiting work or power density.

Although this text has criticised the notion of a power-limit to the energy output of muscle contractions, it was written in undiminished admiration of the groundbreaking work that has been conducted within this conceptual framework. Deciphering the mechanical limits that bind muscle performance across animal size and environments remains challenging enough, and any tool that permits progress should be used. Time only will tell if the kinetic energy density and the physiological similarity index belong into this category, alongside the notion of a power-limit.

## Acknowledgments

This study was supported by a Human Frontier Science Programme Young Investigator Award (RGY0073/2020) to DL and NCH, and partially inspired by an engaging discussion about the Hill- and the Borelli-limit with some members of the Structure and Motion Laboratory at the Royal Veterinary College in London. DL thanks Jim Usherwood for many insightful discussions about muscle work- and power-limits, and for the suggestion to refer to a power-limit as Bennet-Clark’s law. Delyle Polet provided many useful comments on an earlier version of this manuscript, which are gratefully acknowledged.

