## Supplementary Information for "Beyond power limits: the kinetic energy capacity of skeletal muscle"

David Labonte[1] and Natalie C Holt[2]

[1]Department of Bioengineering, Imperial College London, United Kingdom
[2]Department of Evolution, Ecology and Organismal Biology, University of California Riverside, USA

#### Borelli's law and the conservation of energy

Borelli's law is usually presented in terms of a maximum jump height $h$, derived through a balance of the kinetic energy at take-off, and the maximum gravitational potential energy:

$$h = \frac{v^2}{2g} \tag{S 1}$$

Here, $v$ is the take-off speed, and $g$ is the gravitational acceleration; note that this writing implicitly defines the jump height as the height gained after take-off. To predict the variation of jump height with animal size, the take-off speed is then linked with the muscle work capacity $W_{max}$ to find:

$$v = \sqrt{2\frac{W_{max}}{m}} \tag{S 2}$$

where $m$ is the body mass. Animals small and large are assumed to share the same work density, $W_\rho \propto W_{max} m^{-1}$, and consequently are expected to achieve equal take-off velocities, and equal jump heights—Borelli's law:

$$h = \frac{W_{max}}{mg} \tag{S 3}$$

However, this result is only approximate, and in fact violates conservation of energy (Scholz et al., 2006). To understand this objection, note that animals need to lift their centre-of-mass against gravity already during the acceleration phase, i. e., prior to take-off. As a consequence, only part of the muscle work capacity flows into kinetic energy, but this work loss is omitted in the implicit balance between muscle work capacity and gravitational potential energy that underpins the derivation of Borelli's law presented above (Scholz et al., 2006). For small animals, this error is small enough to be negligible, because only a small amount of muscle work output is lost to gravitational potential energy; the work loss does however become meaningful for large animals (Labonte, 2023; Scholz et al., 2006). Thus, strictly speaking, Borelli's law predicts that animals large and small should jump to *approximately* the same height. Even in this approximate form, Borelli's law only holds until animals exceed a critical size; the fraction of work that flows into gravitational potential energy is directly related to the ratio between the gravitational and the ground reaction force, which approaches unity in the largest animals (Labonte, 2023; Labonte et al., 2024).

#### Auxiliary variables and the hidden determination of centre-of-mass dynamics

The dimensional relations between energy, maximum power, speed, mass, and the three auxiliary variables time, force, and displacement, can also be derived from Newtonian mechanics, where they follow quickly from conservation of energy. If the displacement $\delta$ is the variable of choice, conservation of energy dictates $F\delta \propto mv^2$; dividing by $\delta$ and multiplying both sides with $v$ yields $v \propto (2P\delta m^{-1})^{1/3}$. If the time $t$ is chosen, dividing $W \propto mv^2$ by $t$ yields $v \propto \sqrt{Ptm^{-1}}$. If the choice falls on the force $F$, dividing $W \propto F\delta$ by $t$ yields $P \propto Fv$.

In the main manuscript, it is demonstrated that the choice of auxiliary variable uniquely defines the remaining dynamic variables (see Tab. 1 in the main text). These scaling relationships can all be derived in the same way, illustrated here for the case where $P$, $m$ and $\delta$ are fixed, and $v$ is sought. Dimensional consistency demands:

$$[v] = [P]^a[\delta]^b[m]^c$$
$$[\mathrm{L\,T^{-1}}] = [\mathrm{M\,L^2T^{-3}}][\mathrm{L}]^b[\mathrm{M}]^c \tag{S 4}$$

where $a$, $b$ and $c$ are dimensionless numbers, which can be determined by solving a system of simultaneous equations. Matching terms with the dimension mass yields $0 = a + c$; matching terms with the dimension length implies $1 = 2a + b$; and matching terms with the dimension time provides $-1 = -3a$. It thus follows that $a = b = 1/3$ and $c = -1/3$, and thus $v \propto (P\delta m^{-1})^{1/3}$. The scaling relations for all other variable combinations may be derived in the same way.

**What limits contraction time?**

Historically and reasonably, the maximum amount of energy muscle can inject is typically assessed via the path-integral of Newton's second law, i. e., the conservation of energy:

$$F = m\frac{dv}{dt}$$
$$\int F dx = m\int \frac{dv}{dt}dx = m\int v dv$$
$$W = \hat{F}\delta = \Delta E \tag{S 5}$$

where the writing $\hat{F}$ indicates strictly a *displacement-averaged* force. In the main manuscript, it is shown that coupling the conservation of energy with constraints imposed by muscle physiology provides two independent limits to muscle energy output—the Hill-limit, $E = K_{\max} = 1/2mv_{\max}^2$, dictated by the muscle's kinetic energy capacity, and the Borelli-limit, $E = W_{\max} = V\hat{\sigma}\varepsilon_{\max}$, imposed by the muscle's work capacity. Here, we briefly demonstrate that the same result can be found by investigating limits to contraction time, an approach more closely aligned with the premise of Bennet-Clark's law.

The energy output may be written in terms of the contraction time, assumed to be no larger than $t_c$:

$$W = \int F dx \frac{dt}{dt} = \int F v(t) dt = \int^{t_c} \frac{F^2}{m} t dt = \frac{1}{2}\frac{F^2}{m}t_c^2 \tag{S 6}$$

where we used $v = dx/dt = Fm^{-1}t_c$, and assumed for simplicity that the force is constant throughout the contraction. The next step is crucial: what limits the contraction time $t_c$? One possible answer is to borrow a characteristic time from elsewhere, for example by invoking a characteristic frequency through mechanical analyses (e. g. Thangal and Donelan, 2020; Usherwood, 2013; Usherwood and Gladman, 2020), or via a neuro-physiological or sensorimotor control framework (e. g. More and Donelan, 2018; More et al., 2010). Where such a time-scale exists, the muscle energy output may be said to be limited by the muscle's *impulse capacity*, $I = \int F dt$ (Adamson and Whitney, 1971; Knudson, 2009; Ruddock and Winter, 2015; Winter, 2005; Winter et al., 2016). More commonly, however, the time limit is recast in terms of a constraint on displacement (e. g. Bennet-Clark, 1977; Biewener and Patek, 2018; Bobbert, 2013; Gabriel, 1984; James et al., 2007; Marsh, 1994; Sutton et al., 2016). Muscle cannot contract by more than its maximum shortening distance, so that $t_c = \sqrt{2\delta_{max}mF^{-1}}$, and thus $W = F\delta_{max}$—the result is the Borelli-limit, $E = W_{\max}$. However, muscle also cannot contract faster than with an maximum absolute shortening speed $v_{max}$. Reaching this speed takes a time $t_c = v_{max}mF^{-1}$, so that $E = 1/2mv_{max}^2 = K_{\max}$—the Hill-limit (Labonte, 2023).

### Dynamic mechanical performance of a Hill-muscle

For a Hill-type muscle contracting against a payload of mass $m$, the equation of motion reads (Labonte, 2023; Labonte et al., 2024):

$$m\frac{dv}{dt} = F_{\text{max}}\left(\frac{1-v_{\text{rel}}}{1+Qv_{\text{rel}}}\right) \tag{S 7}$$

where we neglected force-length properties and assumed maximum activation for simplicity (a treatment including force-length properties can be found in Labonte, 2023); $F_{\text{max}}$ is the maximum isometric force, $v_{\text{rel}} = vv_{\text{max}}^{-1} \leq 1$ is the relative shortening velocity, and $Q$ is a dimensionless constant, typically of order unity. Where the aim is to assess the energy output of a muscle constrained by a maximum displacement and shortening speed capacity, it is convenient to conduct a variable transformation:

$$m\int v dv = \int F_{\text{max}}\left(\frac{1-v_{\text{rel}}}{1+Qv_{\text{rel}}}\right) d\delta \tag{S 8}$$

This differential equation—a re-expression of Newton's second law as a path-integral—can be solved for any value of $Q$ via separation of variables. However, for all but the case of a linear force velocity relationship ($Q = 0$), the solution does not bring much further joy, and it is thus more convenient to proceed numerically. For $Q = 0$, however, explicit symbolic expressions for several key quantities can be found.

A curious mathematical feature of any Hill-type muscle is that the maximum strain rate is a theoretical construct, only attainable in the the limit of an infinitely long contraction. No contraction can be infinitely long however, so that a Hill-type muscle always reaches its displacement capacity before it is contracting with its maximum strain rate (see Labonte, 2023; Labonte et al., 2024, and below). The shortening speed that arises from a contraction over the full displacement capacity then follows by fixing the integration limit of the displacement integral in eq. S 8 to $\delta_{\text{max}}$ (Labonte, 2023; Labonte et al., 2024):

$$v_{\text{Hi-Bo}} = v_{\text{max}}\left[1+\text{W}\left(-\exp\left(-1-\frac{1}{2\Gamma}\right)\right)\right] \tag{S 9}$$

where W is the Lambert W (or ProductLog) function, and $\Gamma = K_{\text{max}}/W_{\text{max}}$ is the physiological similarity index—the ratio between the maximum kinetic energy and work capacity for a muscle for which the force-velocity-relationship is a simple step-function (Labonte, 2023; Labonte et al., 2024). The work done by the muscle then readily follows by recognising that $E = 1/2mv_{\text{Hi-Bo}}^2$, so that:

$$W_{\text{Hi-Bo}} = K_{\text{max}}\left[1+\text{W}\left(-\exp\left(-1-\frac{1}{2\Gamma}\right)\right)\right]^2 \tag{S 10}$$

The kinetic energy capacity is always the same, no matter the specific form of the FVR, so that $\Gamma_{\text{Hi-Bo}}$ is $\Gamma_{\text{Hi-Bo}} = K_{max}/W_{\text{Hi-Bo}}$, or:

$$\Gamma_{\text{Hi-Bo}} = \left[1+W\left[-\exp\left(-1-\frac{1}{2\Gamma}\right)\right]\right]^{-2} \tag{S 11}$$

The displacement-averaged force, in turn, is $\hat{F}_{\text{Hi-Bo}} = W_{\text{Hi-Bo}}/\delta_{\text{max}}$, and thus:

$$\hat{F}_{\text{Hi-Bo}} = \hat{F}_{\text{K}}\left[1+\text{W}\left(-\exp\left(-1-\frac{1}{2\Gamma}\right)\right)\right]^2 \tag{S 12}$$

where $\hat{F}_{\text{K}}$ is the average inertial force, $K_{max}/\delta_{\text{max}}$. This completes the derivation of the three results leaned upon in the main manuscript (eqs. 5-7).

**Is real muscle ever limited by $K_{\text{max}}$?**

In the main manuscript, it is concluded that the energy output of a musculoskeletal system is limited by its kinetic energy capacity $\Gamma \leq 1$, and by its work capacity for $\Gamma \geq 1$. For a Hill-type muscle, it was derived above that:

$$\Gamma_{\text{Hi-Bo}} = \left[1 + W\left[-\exp\left(-1 - \frac{1}{2\Gamma}\right)\right]\right]^2 \tag{S 13}$$

where $\Gamma$ is the physiological similarity index for a muscle that has a force-velocity-relationship idealised as a step function:

$$\Gamma = \frac{1}{2}\frac{m}{V} l_m^2 \frac{\dot{\varepsilon}_{\text{max}}^2}{\sigma_{\text{max}} \varepsilon_{\text{max}}} \tag{S 14}$$

Equation S 13 is not particularly forthcoming, but of key relevance are its limits: for $\Gamma \to 0$ and $\Gamma \to \infty$, one has $\Gamma_{\text{Hi-Bo}} \to 1$ and $\Gamma_{\text{Hi-Bo}} \to \infty$, respectively. In other words, $\Gamma > 0$ but $\Gamma_{\text{Hi-Bo}} > 1$ (Fig. S 1). The energy output of a Hill-type muscle may thus get arbitrarily close to its kinetic energy capacity, but it will notationally always be displacement- and thus Borelli-limited—but this is strictly a purely mathematical and not a physiological or physical result (see also the SI in Labonte, 2023; Labonte et al., 2024). The physical implication is instead that there exist two equally valid interpretations for what limits the energy output at small $\Gamma$: the work capacity is reduced far below the quasi-static maximum by the variation of stress with strain rate (Bobbert, 2013), or the muscle's kinetic energy capacity is insufficient. Which interpretation is preferred is to some extent a philosophical decision. However, one may argue that it remains advantageous to define the physiological similarity index via eq. S 14 instead of via eq. S 13—and to thus conclude that the kinetic energy capacity is limiting—for at least three reasons. First, eq. S 14 is easy to evaluate analytically, does not depend on the exact form of the Hill-relation, and retains the correct asymptotic limits. Second, $\Gamma_{\text{Hi-Bo}}$ varies only little for small $\Gamma$, although the associated mechanical output can vary quite substantially (see main manuscript). And third, it provides the opportunity to distinguish between two distinct effects of any FVR: the maximum strain rate as such, the variation of stress with strain rate.

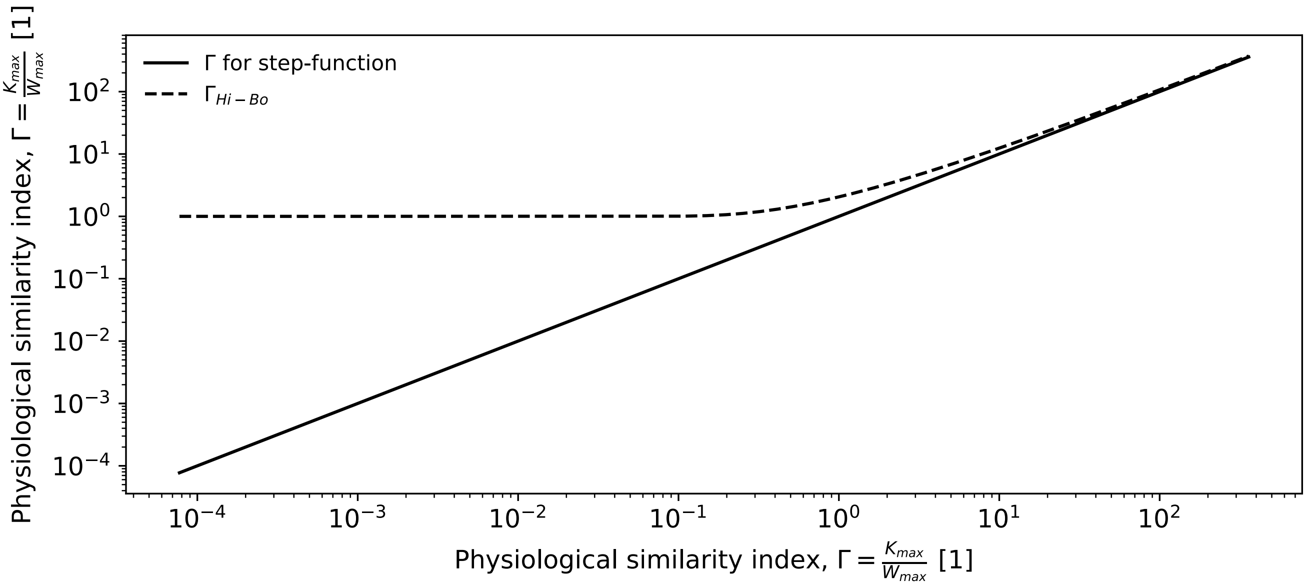


Figure S 1 | Comparison of the physiological similarity index for a muscle that has a force-velocity-relationship idealised as a step-function, vs a muscle that has a linear force-velocity relationship. For the former, $\Gamma > 0$, but for the latter, $\Gamma_{\text{Hi-Bo}} > 1$—a Hill-type muscle is thus notationally always displacement-limited. However, its energy capacity approaches $K_{\text{max}}$ with arbitrary accuracy, so that the distinction between the kinetic energy and maximum work capacity remains insightful and useful.
